# Molecular goniometers for single-particle cryo-EM of DNA-binding proteins

**DOI:** 10.1101/2020.02.27.968883

**Authors:** Tural Aksel, Zanlin Yu, Yifan Cheng, Shawn M. Douglas

## Abstract

Correct reconstruction of macromolecular structure by cryo-electron microscopy relies on accurate determination of the orientation of single-particle images. For small (<100 kDa) DNA-binding proteins, obtaining particle images with sufficiently asymmetric features to correctly guide alignment is challenging. DNA nanotechnology was conceived as a potential tool for building host nanostructures to prescribe the locations and orientations of docked proteins. We used DNA origami to construct molecular goniometers—instruments to precisely orient objects—to dock a DNA-binding protein on a double-helix stage that has user-programmable tilt and rotation angles. Each protein orientation maps to a distinct barcode pattern specifying particle classification and angle assignment. We used goniometers to obtain a 6.5 Å structure of BurrH, an 82-kDa DNA-binding protein whose helical pseudosymmetry prevents accurate image orientation using classical cryo-EM. Our approach should be adaptable for other DNA-binding proteins, and a wide variety of other small proteins, by fusing DNA binding domains to them.

---

Single-particle cryo-electron microscopy (cryo-EM) has recently emerged as a powerful technique for achieving high-resolution structure determination of biological molecules^1,2^. A typical cryo-EM study includes adsorbing a specimen onto a grid, freezing the specimen within a thin layer of vitreous ice, collecting cryo-EM images of individual particles, computationally sorting and aligning the particle images, and reconstructing a 3D structure from 2D particle images. In the final step, an algorithm refines orientations of the 2D particles and determines the 3D reconstruction via iterative expectation-maximization optimization. Refinements of the 3D reconstruction and 2D particle orientations typically converge to a correct solution when particle image data are sufficient in quantity, resolution, distribution of orientations, and conformational homogeneity. However, even if these conditions are met, when particles lack obvious asymmetric features to guide the alignment—as is often the case for small <100 kDa targets—the optimization gets trapped in a local minimum and thus yields a low-resolution model. Others have sought to overcome this problem by, for example, docking the targets to protein scaffolds such as antibody fragments^3^, bacterial ribosomes^4^, or engineered nanoparticles^5^. Scaffold-based approaches typically rely on rigid attachment of target proteins to the scaffold, and the structure of the full complex is determined all at once. However, no one has yet been able to generalize these solutions and successfully use them as routes to determining structures of classes of targets where a rigid attachment is impractical.

Ned Seeman envisioned programming DNA, via self-assembly, into custom nanostructures as a way to engineer positional control of biomolecules for crystallographic studies of proteins^6^. Nearly 40 years later, DNA nanotechnology can now enable the construction of tiny mechanical devices, which offer opportunities to reconceptualize certain challenges when visualizing structures of biomolecules^7^. In 2006, Rothemund introduced DNA origami, a powerful approach to building megadalton-sized nanostructures that relies on a long single-strand DNA (ssDNA) ‘scaffold’ to template the assembly of numerous short DNA oligonucleotide ‘staples’ into a custom shape^8^. This method has since been used to build several innovative custom instruments helpful for measuring and identifying forces in biological systems^9–12^. As a way to address an important challenge of cryo-EM—difficulty in aligning small particle images—we aimed to correctly determine each 2D particle orientation by *prescribing* the orientation using a nanoscale device: a molecular goniometer made out of DNA origami. Our work builds on ideas developed by Dietz, Scheres, and colleagues^13^, and includes several optimizations that helped improve resolution of 3D reconstructions generated with their related approach.

We describe the design, characterization, and validation of molecular goniometers. Since the different orientations of small protein images are often too subtle to determine directly, we mapped unique angle configurations to large asymmetric features, or barcodes, on the DNA origami. We validated our approach by obtaining a 6.5 Å structure of BurrH^14^, an 82 kDa TALE-family DNA-binding protein whose crystal structure is known (PDB id: 4cja). We found that 3D structure maps built without the *a priori* information provided by the DNA origami (i.e., the tilt and rotation angle “priors”) did not fit as well to the known structure as the final map we constructed using the priors.

## Results

Our molecular goniometers enable user-controlled tilt and rotation of single proteins, coupled with large asymmetric features (barcode “bits”) to provide unambiguous identification of each protein orientation. We demonstrate the feasibility of determining the structure of a small protein attached to a host nanostructure by cropping out (or masking) the host particle and using only the subimages containing each protein together with its orientation information obtained from the host.

Our design includes key conceptual and technical advances. One important advance our work achieved is that we show that a perfectly rigid link between the protein and the host nanostructure is not required, and may not be desired, for this style of structure determination. In fact, we found it useful to isolate and separately tune *spatial* and *angular* rigidity. Spatial rigidity is desirable when positioning the protein for well-isolated, well-centered extraction of low-background subimages, but care must be taken with the design as increasing separation between the protein and host can compromise rigidity, and vice versa. Angular rigidity between the protein and the goniometer is also needed, otherwise the angle priors do not have a meaningful relationship with the true orientation of the protein. On the other hand, excess angular rigidity may limit the particle-angle distribution and compromise reconstruction resolution. Therefore, our design allows for limited swiveling of the protein rotation angle, centered around a target angle. Finally, the goniometers must be designed to self-assemble with a high-enough yield to maintain precision and accuracy while satisfying these constraints.

Figure 1 summarizes the molecular goniometer design, which consists of a fixed chassis (gray), a programmable DNA stage (yellow) containing a protein-binding site (magenta), and barcode domains (teal) that uniquely identify each stage-angle configuration. The chassis resembles a tiny C-clamp that grasps the DNA stage, which is a 56-base-pair (bp) double helix anchored on both sides of the chassis aperture. The DNA stage orientation is set by the staple sequences in the anchoring regions of the chassis (dark gray) and by the 5’-to-3’ direction of the scaffold sequence within the origami.

**Fig. 1.**
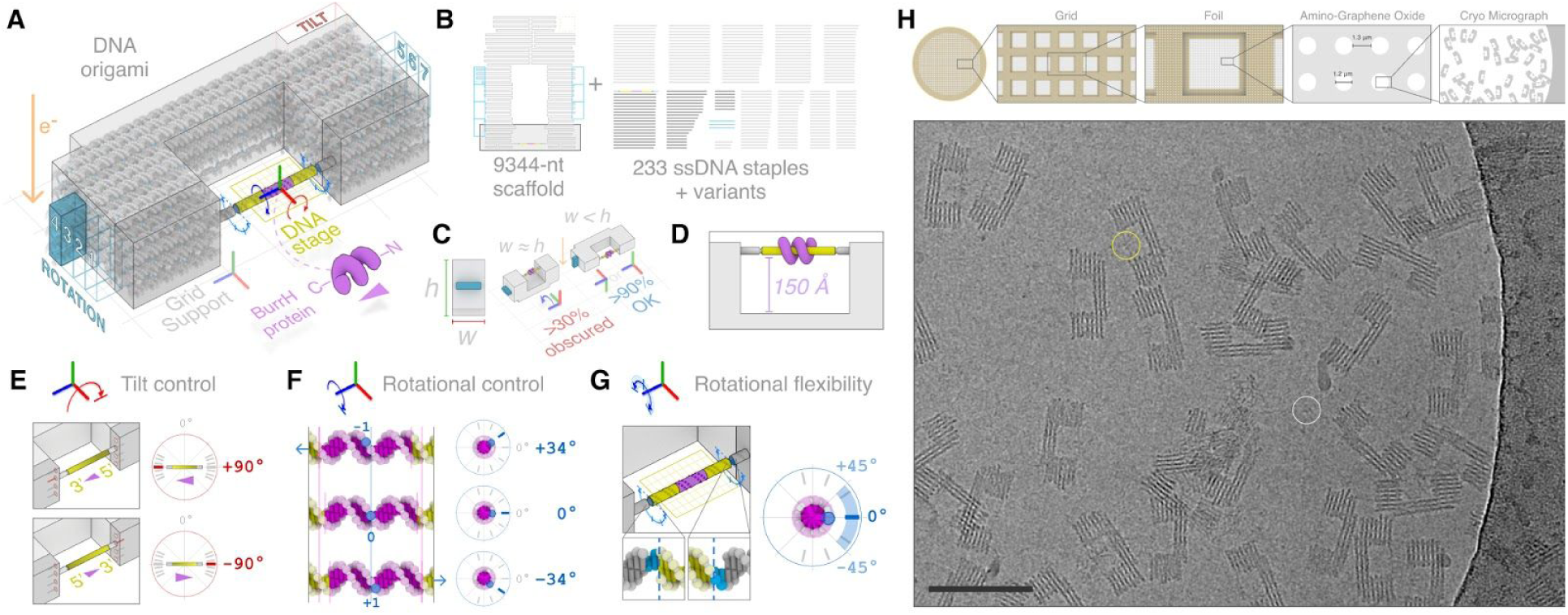
Design and grid adsorption of molecular goniometers built from DNA origami. (**A**) A representative configuration is shown (+90° tilt, 0° rotation). DNA origami chassis (grey) grasps and controls the tilt and rotation of the 56-bp DNA stage (yellow) containing the 19-bp binding site for the BurrH protein (magenta). The TILT barcode bit (salmon) and ROTATION barcode domains (teal) identify each unique DNA stage configuration. (**B**) 2D schematic of origami scaffold and staples. (**C**) Chassis aspect ratio can promote the preferred orientation of goniometer adsorption onto the grid surface. (**D**) The DNA stage is positioned with a 150 Å gap from the origami chassis. (**E**) DNA stage tilt angle is set by the polarity of the origami scaffold route and the chassis attachment locations. (**F**) Rotation angle is controlled by shifting the register of the DNA stage relative to the chassis. A 1-nt linear shift of the stage corresponds to a 34° angular rotation of the protein binding site. (**G**) The DNA stage is flanked by unpaired 2-nt regions to provide rotational flexibility. An SD of 15° is used for the Gaussian rotation priors. (H) A mixture containing BurrH and goniometers (at a 10:1 molar ratio) is deposited on a gold quantifoil grid with an amino graphene oxide support. A representative cropped micrograph is shown. Circles indicate typical bound BurrH (yellow circle) and unbound BurrH (white circle). Scale bar: 100 nm.

The DNA stage (yellow) includes the 19-bp binding sequence of the BurrH DNA-binding protein (magenta). The outer edges of the goniometer chassis can be configured to display modular barcode domains, or “bits”, which form unique asymmetric patterns that enable particle classification. Our barcode design, which we expand upon below, includes seven bits (1–7) that specify the rotation angle of the DNA stage, and one bit (labeled ‘TILT’) that specifies the tilt angle of the DNA stage. The barcode pattern is set by including the corresponding ssDNA staples in the origami folding reaction. In Fig. 1A, the 4th bit of the rotation barcode is enabled, corresponding to a stage rotation angle of 0°. The remaining disabled rotation bits (1–3, 5–7) are shown as teal outlines. The location of the disabled tilt bit is outlined in salmon red.

The molecular goniometer is comprised of a 9344 nucleotide (nt) scaffold and 233 staples (strand diagram, Fig. 1B). Colors of boxed scaffold regions and staples indicate the corresponding features from Fig. 1A. The DNA stage is comprised of a cloned scaffold segment and a staple strand (Fig. S1). The scaffold arrangement in the dark gray zone is modified, along with corresponding barcode regions, to change the angle settings.

To ensure that the molecular goniometer chassis adsorbs onto the grid in a desired orientation relative to the electron beam direction, we designed the chassis cross-section to have a narrow aspect ratio (Fig. 1C). We found that a cross-sectional aspect ratio of w/h < 0.6 helped approximately 90% of the goniometers adsorb to the grid surface in a preferred orientation (face-up or face-down), positioning the DNA stage without obscuring the protein (Fig. S2).

To minimize protein image artifacts due to delocalized signal from the origami, we positioned the DNA stage to ensure a large (150 Å) gap between the protein and chassis (Fig. 1D). Smaller than that, Fresnel fringes originating from the origami will overlap with the protein particle image, interfering with isolating the protein particles for image processing.

The tilt angle of the DNA stage is controlled by the anchor locations on the chassis, and by the polarity of the scaffold strand (Fig. 1E). Rerouting the scaffold onto the complementary strand within the origami design, in the opposite 5’-to-3’ direction relative to the chassis, flips the stage tilt angle by from +90° to –90°. Therefore, goniometer designs with the +90 and –90° tilt angles each require a separate set of staple strands.

The rotation angle of the protein-binding site is controlled by the register of the stage sequence relative to the goniometer chassis (Fig. 1F). Assuming 10.5 bp per helical turn, a 1-nt shift corresponds to a 3.4 Å translation along the stage axis, and a 34.3° rotation. To modify the stage rotation angle, we can modify the scaffold route in the surrounding anchoring region (dark gray, Figs. 1A & 1B), with corresponding updates to the complementary staple sequences (Fig. 1B, dark gray). We designed goniometers with seven possible rotation angles, and assigned the rotation angle 0° to the middle angle. Thus, the possible rotation angles, rounded to the nearest integer, are +103°, +69°, +34°, 0°, –34°, –69°, and –103°. Figure S3 explains how we define the goniometer tilt and rotation angles.

Each tilt and rotation angle configuration is paired with a unique barcode pattern that is similarly implemented via local modifications to the scaffold route and corresponding staple sequences. The 1-nt resolution of DNA stage registration corresponds to a rotation angle step size of ∼34° which, despite some intrinsic torsional flexibility, may not provide sufficient angle coverage for cryo-EM structure determination. To ensure sufficient variance in the distribution of rotation angles, the DNA stage region is flanked on both sides by 2-nt ssDNA regions (blue), as shown in Fig. 1G.

A pooled mixture with equal fractions of all goniometer designs are adsorbed onto holey gold grids with an amino-graphene oxide support^15^; Fig. 1H shows a representative micrograph region. We collected a total of 17,725 images. We were able to identify 908k goniometer particles. Of those, 669k (74%) had identifiable tilt and rotation barcodes, 398k (44%) had BurrH present. We selected 211k BurrH particles with well-folded protein and low background signal. Thus, 23% of all goniometers had good BurrH particles present, and representing an average of 12 good BurrH particles per image.

Figure 2 describes how we classified the goniometers by their barcodes and then assigned tilt- and rotation-angle priors to each masked protein particle, which are used as inputs for building the initial 3D reconstruction. After cryo-EM micrograph data collection, particles are processed in a multi-step computational pipeline (Fig 2A). Individual goniometer particles are picked and aligned into classes, which are then sorted into +90° and –90° tilt-angle classes according to the tilt-angle barcode domain (Fig. 2B).

**Fig. 2.**
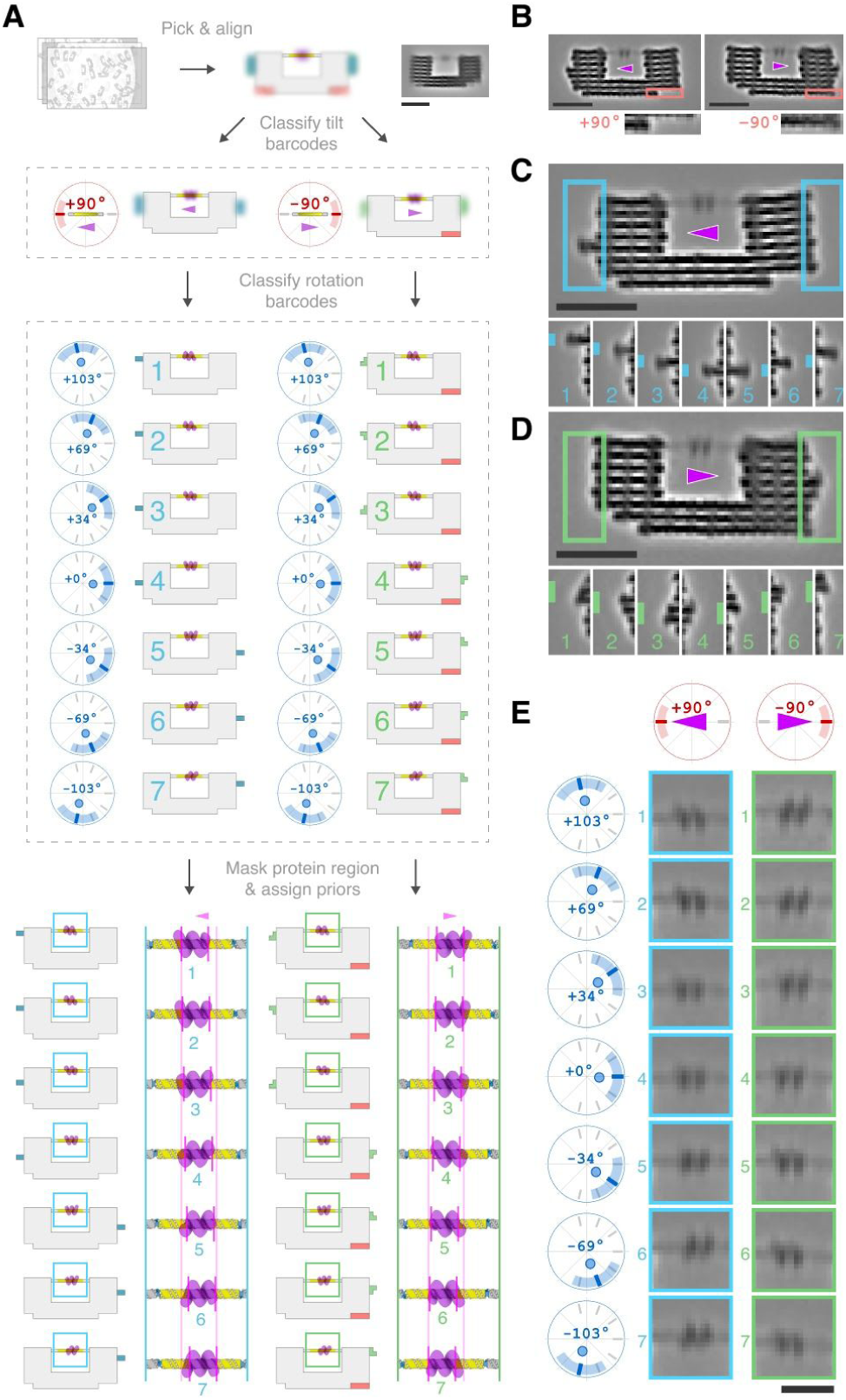
Determination of protein classes with angle priors from goniometer image data. (**A**) Goniometer data processing pipeline. Top right: representative 2D class, prior to sorting by tilt barcode. (**B**) Example +90° and –90° tilt classes, with barcode detail below. Only the –90° tilt class has the 582-nt barcode bit enabled. Magenta arrowhead indicates N-to-C orientation of BurrH. (**C**) The 222-nt rotation barcode bit is comprised of 108-nt scaffold paired with 3 staples of 114-nt combined length. (**D**) An alternate 414-nt bit design (186 nt scaffold + 228 nt combined staple length) was also used. (**E**) 2D consensus classes derived from goniometers with stages configured at +90° and –90° tilt angles, and 7 rotation angles (0°, ±34°, ±69°, and ±103°). Scale bars: (A–D) 20 nm, (E) 10 nm.

The differences between each DNA stage configuration would be difficult to classify directly, so we rely on decorating the goniometers with machine-readable geometric patterns, or barcodes, that identify the DNA stage tilt and rotation configuration. Each barcode pattern is comprised of two bits that can be toggled on (or off) by routing (or not routing) the scaffold and staples to fold into small domains the edges of the chassis.

We used a single 582-nt bit (Fig. 2B) to identify frames with a –90° tilt angle. The tilt subclasses were further divided into groups by rotation-angle barcodes. We tested two rotation-angle barcode designs: a 222-nt bit (Fig. 2C), and a 414-nt bit (Fig. 2D). The 414-nt bit was used with both +90 and –90 tilt designs, while the 222-nt bit was used only for +90 tilt (Fig. S2), and data from all goniometer variants were combined for the final structure. Of the two barcode designs that we tried, the smaller rotation-angle barcode bit (Fig. 2C) seems to work best for classifying particles, perhaps because it projects farther from the edge of the origami. The 222-nt bit was successfully classified 84% of time, versus 70–75% for the 414-nt bits.

After the goniometers are classified according to tilt and rotation barcodes, a mask is used to isolate the DNA stage and protein from the chassis. We generated 2D consensus classes for each angle configuration (Fig. 2E), in which different orientations of BurrH are clearly visible. Altogether, we tested 21 different goniometers (Fig. S4). Seven goniometers used a +90° tilt angle (with the tilt bit disabled), and each respective rotation angle (–103°, –69°, –34°, 0°, 34°, 69°, 103°) identified with a 222-nt bit. Seven goniometers also used a +90° tilt angle and same rotation angles, but with 414-nt rotation bit. Finally, seven goniometers used a –90° tilt angle (tilt bit enabled), and seven rotation angles identified with a 414-nt bit. All goniometers were pooled for the final 3D reconstruction (Fig. S5).

We used Gaussian priors to constrain the assignment of particle angle and position parameters within a window centered on a mean value. Prior to 3D reconstruction, we flipped the BurrH particle images derived from goniometers with –90° tilt so that all BurrH particles have the same +90° tilt angle. To constrain tilt angles for all particles, we used a Gaussian prior centered at +90° with a window size of ±30°. For the rotation angles, we used Gaussian priors centered at the rotation goal angles specific to each goniometer class, with a window size of ±45°. Figure S6 describes the process of barcode classification, and Figure S7 includes twenty 2D class averages for each particle.

Figure 3 shows the final 3D reconstruction of BurrH using the *a priori* orientational information derived from our molecular goniometers (for detail, see Fig. S8), along with an assessment of the tilt and rotation angle accuracy, and a side-by-side comparison with 3D reconstructions built without and with the angle priors.

**Fig. 3.**
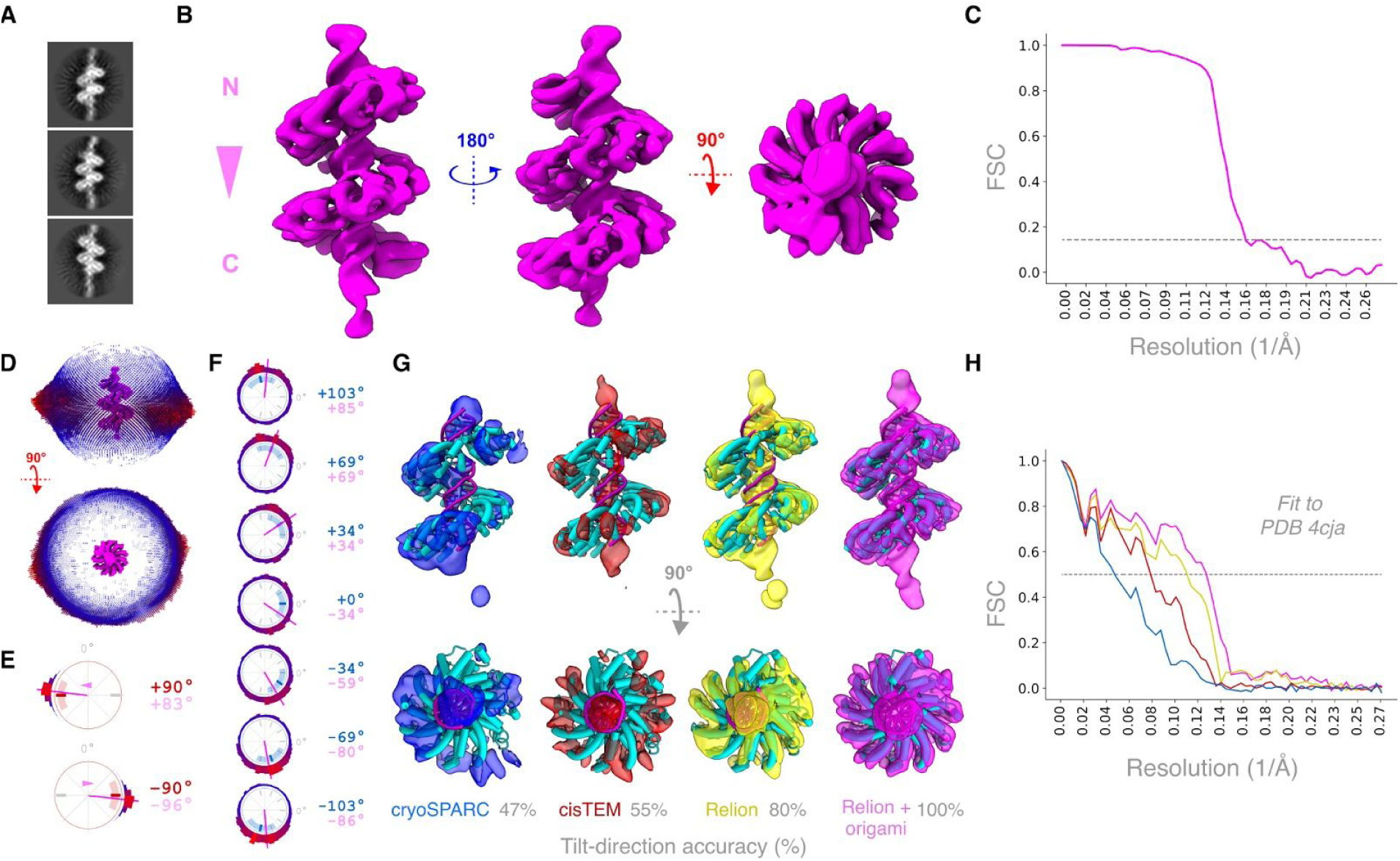
3D refinement of BurrH structure using molecular goniometer-derived angle priors. (**A**) Representative BurrH 2D class averages. (**B**) 3D reconstruction of BurrH using *a priori* tilt and rotation angles from molecular goniometers. (**C**) FSC curve calculated from two half-maps of the final 3D reconstruction. FSC_0.143_ line crosses the curve at 6.5 Å resolution. (**D**) Euler angle distribution derived from 3D refinement, with blue-red color scale. (**E**) Representative polar plots for stage tilt angles for +90° and –90° designs. The inner red tick mark represents the goal tilt angle, and the ±45° arc shows the bounds of the Gaussian tilt angle prior. The measured tilt angle distributions are plotted as histograms. Average tilt angles (magenta line) were +83° and –96°, respectively. (**F**) Representative polar angle plots for seven stage rotation angles, derived from goniometers with +90° tilt and 414-nt barcode bits. The inner blue tick mark represents the goal rotation angle (integer value in blue text), and the ±45° arc shows the bounds of the Gaussian rotation angle prior. Average rotation angles are plotted as a magenta lines (integer values in pink text). (**G**) Comparison of 3D reconstructions obtained without angle priors (blue: cryoSPARC, red: cisTEM, yellow: Relion), and with priors (magenta: Relion with goniometer-derived angle priors), with BurrH crystal structure (cyan) fit to each density map. Areas where cyan is visible indicate a poor fit. Tilt-direction accuracy is compared to the true orientations derived from gonimeters. (**H**) FSC curves showing goodness-of-fit of BurrH crystal structure into 3D cryo-EM density maps of BurrH. FSC curve is computed following a rigid body fit of BurrH crystal structure into each cryo-EM density map. FSC_0.5_ shown as dashed line.

To build our 3D reconstruction of BurrH aided by goniometer-derived angle priors, we used the software Relion^16^ and custom Python code, with the following approach: With both tilt- and rotation-angle constraints enabled, we first generated an initial model that became the starting structure for the 3D reconstruction. We then aligned all particles to the initial model with both tilt and rotation angle constraints enabled. Finally, during the subsequent 3D classification of particles and the refinement of the best 3D class, we removed the rotation-angle constraint. We kept the tilt-angle constraint enabled at every step of 3D reconstruction because we were highly confident about the direction of protein particles bound to the stage DNA.

Figure 3A shows representative good class averages from the 3D model (for additional classes, see Fig. S9). The final 3D reconstruction of BurrH (Fig. 3B) has all the repeats and the secondary structures resolved with an estimated resolution of 6.5 Å (Fig. 3B, C). Tilt and rotation angle values from the final refinement are well-distributed with a good coverage of Euler angles (Fig. 3D). The observed angle values were within ±45° of the goal angles for all goniometer designs (Fig. 3E,F). See Fig. S10 for the complete angle distributions of all designs.

To evaluate the utility of our overall approach, we performed three additional 3D reconstructions of BurrH without the use of *a priori* orientation information (i.e., the angle priors) provided by the molecular goniometers. We used cryoSPARC ^17^, cisTEM ^18^, Relion ^16^ for these reconstructions, and then compared the results to our own approach, which used Relion along with the origami-derived priors.

For 3D reconstructions without angle priors, we used the same initial model and the particles from the final refinement step of the 3D reconstruction with the angle priors. In Fig. 3G, the 3D maps were each fit to the PDB structure, and the tilt-direction accuracy for each map was based on a comparison with the true angles derived from the origami particles. In all reconstructions that did not use angle priors, some fraction of particles ranging from 20–53% are misaligned with respect to the stage DNA axis (Fig. 3G). The BurrH crystal structure (Fig. 3G, cyan) does not fit well in the 3D volumes obtained from the reconstructions without angle priors and the correlation between the BurrH crystal structure and the 3D reconstruction volumes gets worse with the fraction of misaligned particles (Fig. 3H).

## Discussion

The success of our approach depends on multiple key features of the goniometers, many of which we can evaluate in the context of a closely related work published in 2016 that described a DNA origami molecular support for protecting proteins from air-water interface exposure during cryo-EM sample preparation^13^. Martin *et al.* used their device to obtain a 15 Å reconstruction of the transcription factor p53—a 160 kDa tetramer—and pioneered several design and analysis concepts that we adopted, including sub-image masking to isolate the protein prior to reconstruction, the general aim of controlled protein orientation. The authors also identified many challenges that limited the initial success of the approach, including a low yield of intact particles, signal delocalization due to imaging defocus, and difficulty in controlling the protein orientation. In our design, we sought to maximize the yield of intact protein-bearing particles per micrograph, especially since the goniometer particles are massive (>6 MDa) and occupy a large footprint on the grid (>2000 nm^2^). We attempted using ssDNA staples to create the DNA stage, but the fraction of goniometers with bound protein was below 10%. Cloning one strand of the DNA stage into the origami scaffold increased the occupancy to 60%, presumably due to enrichment of intact protein-binding substrates (Fig. S1). Adopting a chassis with a narrow aspect ratio boosted the fraction of particles in desirable orientations by 30% relative to a chassis design with a wider aspect ratio (Fig. S2). To mitigate superposition of the signal from the support structure and the target protein, we designed our chassis to position the DNA stage far from the chassis (Fig. 1A, D).

Contrary to what might be expected, our designs intentionally did not maximize the rigidity of the DNA stage with respect to the chassis. We wanted to obtain uniform sampling of rotation angles to avoid compromising the resolution of the 3D reconstruction^19^. Therefore, we included 2-nt gaps flanking the DNA stage to provide rotational flexibility and angle variance. It is unclear why some DNA stage configurations did not average to exactly their goal angles. However, the observed deviations were within tolerable ranges for our 3D reconstruction process. If future studies require greater rotational angle precision, our designs could be modified to reduce one or both of the 2-nt gaps to 1-nt gaps or 0-nt nicks, or enzymatically ligated to further increase rigidity. Additional studies could establish design rules for finely tuning the accuracy and precision of rotation angles.

We tested goniometers designed with +90° and –90° tilt angles, but were unable to resolve any difference between the designs with the amount of data we collected. Since the rotational flexibility seemed to provide a uniform and even sampling of rotation angles, we treated the +90 and –90 tilt angles as equivalent during our analysis, although this assumption may not hold for DNA stages designed with less rotational flexibility. While we only tested two tilt angles, each with seven rotation angles, the goniometer allows for additional angles. The chassis accommodates tilt angles from 65° to 115° (or –65° to –115°), with a 7–10° step-size, based on rerouting the scaffold through alternate DNA stage anchor locations. The DNA stage can be set to 21 distinct rotation values, representing a 17° step size. Further testing will be needed to determine if and when different angle configurations are useful, for example for proteins that exhibit symmetry along a different axes compared to BurrH.

We compared how three popular cryo-EM software tools would handle determination of the 3D model and orientations of the 2D particles. We offer this comparison as a practical guide for cases where angle priors are necessary (i.e., when particles lack asymmetric features to guide alignment). The tools cryoSPARC and cisTEM do not accept user-specified Gaussian angle priors as input, so we were unable to test the quality of model they might build with that information. Moreover, the comparison may not reflect how the tools would perform given particles that had not already been picked, aligned, and masked from the DNA origami goniometers, which effectively serve as fiducial markers in the first step of image analysis. However, starting with the centered and masked BurrH particles allows for a well-controlled comparison of how the tools handle angle assignment specifically. Although Relion and cisTEM both utilize a maximum-likelihood algorithm, Relion achieves greater accuracy (80%) without priors compared to cisTEM (55%). CryoSPARC uses a conjugate-gradient optimization which was the fastest, but also the least accurate (47%). Relion performed better than random in determining the correct tilt angles without any priors, perhaps due to performing a more extensive search of the optimization space. However, Relion was incorrect for enough particles (20%) that it could not match the resolution compared to using the priors.

Our final reconstruction of BurrH achieved a resolution of 6.5 Å using 68,482 particles. We collected micrographs at 22,000× magnification, corresponding to a pixel size of 1.82 Å, and maximum resolution of 3.64 Å. BurrH was expected to be a challenging target due to its small size and helical pseudosymmetry, but additional factors may further complicate its structure determination with cryo-EM. If flexibility of the DNA stage-BurrH complex resulted in some conformational heterogeneity of our dataset, then collecting additional images may drive the resolution higher. It is also possible that the current generation of computational tools has difficulty distinguishing dynamic variability in small structures; further algorithmic optimizations may be beneficial. Because we can selectively modify several design parameters, our goniometers may be useful for systematic and well-controlled testing of the limits of experimental and computational methods in cryo-EM studies of small proteins.

We obtained an average of 51 goniometers per micrograph, 24% of which had a “good” BurrH used in the initial reconstruction. The overall goniometer particle density on the grid surface might be increased by reducing the mass and footprint of the origami by using a more compact design, and promoting uniform orientation of chassis adsorption onto the support surface, for example, using a custom affinity grid and staples with anchoring ssDNAs that extend from one face of the goniometer. We can also improve the capacity for sample multiplexing on a single grid by using more complex barcodes.

Of the six degrees of freedom (x, y, z, tilt, rotation, psi) that specify the absolute location of a particle on a cryo-EM grid, our molecular goniometers provide limited user-defined control for three of them (tilt, rotation, and z translation from the grid surface). We designed the DNA origami particles to be large, asymmetric, and easy to pick and align, and so in effect they do provide local x, y, and psi values of the DNA stage relative to the origami chassis. The capability of prescribing the distance of the target from the surface of the grid could be useful for certain targets, perhaps including targets with masses greater than 100 kDa with some modifications to the origami design. It should also be possible to gain better control over additional degrees of freedom, such as the absolute x, y, and psi values.

In cases where a tool or algorithm could achieve 100% tilt angle accuracy without the aid of priors derived from the origami, then the goniometers would not be necessary. However, currently this level of accuracy is not always possible. Thus, our results demonstrate that the *a priori* information provided by the molecular goniometers improve resolution for cryo-EM studies of small DNA-binding proteins such as BurrH. Furthermore, the paucity of small (<100 kDa) proteins in the Electron Microscopy Data Bank may suggest that there are many targets of unknown structure that may benefit from our approach, either by incorporating their binding sequence into the DNA stage, or for non-DNA-binding proteins, fusion with a DNA-binding domain. In principle, RNAs can bind to the DNA stage directly via hybridization, or by docking to RNA-binding moieties that attach to the DNA stage.

Finally, while the field of cryo-EM has seen great technical advancements in recent years, many areas could be improved and significant limitations to its use exist ^20,21^. Using nanotechnology to take control of the positions and orientations of the particles themselves may provide a uniquely powerful tool in the effort to bridge the gap between the practical and theoretical limits of the method.

## Acknowledgments

We thank F. Wang for furnishing amino graphene oxide grids for early tests. We thank J. Brown and C. Gingold for help with visualization.

## Author contributions

TA and SMD conceptualized the project. TA performed research, collected data, wrote software, and analyzed data. ZY performed research and collected data. All authors discussed data. TA and SMD wrote the manuscript with input from all authors. YC and SMD provided resources and supervised the project

## Data and materials availability

Source code is available at https://github.com/douglaslab/cryoorigami. The cryo-EM map of BurrH has been deposited in the Electron Microscopy Data Bank (EMD-21443). The cryo-EM movie files have been deposited at the Electron Microscopy Public Image Archive (EMPIAR-10373). The p9344-BurrH scaffold sequence has been submitted to GenBank (MT081208), and the plasmid deposited with AddGene (#140326).

## Supplementary Materials

### Materials and Methods

#### P rotein Expression and Purification

The BurrH open reading frame (ORF) with a flexible linker and C-terminal HIS TAG (-GSGHHHHHH) was subcloned into pet24d+ expression vector (Genscript, NJ, USA). For protein concentration quantification, we added single tryptophan residue upstream of the HIS TAG. The BurrH plasmid was chemically transformed into *E.coli* strain BL21(DE3)pLysS (Promega, WI, USA) as described in the product manual. BurrH was expressed and purified as described^22^. Briefly,

1. 10 ml sterile LB media supplemented with 100 µg/ml Ampicillin was inoculated with a single colony picked from an LB-agarose plate. The bacterial culture was grown overnight at 37°C with moderate shaking.
2. The next day, 10 ml of the overnight culture was added into 1L sterile LB media supplemented with 100 µg/ml Ampicillin and the 1L culture was grown at 37°C with moderate shaking for 4–6 hours until OD600 reaches 0.4–0.6.
3. Once OD600 reaches 0.4–0.6, culture was brought to 1mM IPTG with a sterile 1M IPTG stock. The culture was grown at 37°C with moderate shaking for an additional 4 hours.
4. Cells were harvested by centrifuging the culture at 6000× rcf for 10 minutes at 4°C.
5. For long-term storage and easy lysis of E.coli cells, the cell pellets were kept at −80°C.
6. Cell pellets were lysed using B-PER Bacterial Protein Extraction Reagent (ThermoFisher Scientific, MA, USA). 4 ml B-PER reagent was added per gram of cell pellet. In addition, the lysis reagent was supplemented with 100 µg/ml lysozyme and 3U DNase I. The lysate was incubated at room temperature with moderate stirring for 15 minutes.
7. Lysate was centrifuged at 15,000× rcf for 15 minutes and the pellet was discarded.
8. Clear lysate was filtered using a 0.45-µm syringe filter (Millipore Sigma, MA, USA).
9. Lysate was mixed with 2 ml nickel NTA beads (Qiagen, MD, USA) equilibrated with wash solution (10% Glycerol, 150 mM NaCl, 20 mM Imidazole, 50 mM HEPES-Cl at pH 8.0). The lysate-and-nickel-beads mix was incubated at 4°C with moderate mixing for 30 minutes.
10. Beads were washed twice using 25 ml wash solution each time.
11. BurrH was eluted using 10 ml elution solution (10% Glycerol, 150 mM NaCl, 250 mM Imidazole, 50 mM HEPES-Cl at pH 8.0) and passed through a 10-ml Pierce disposable column (ThermoFisher Scientific, MA, USA) to remove the beads from solution.
12. The final elution was dialyzed against BurrH solution (10 mM MgCl2, 150 mM NaCl, 50 mM HEPES-Cl at pH 8.0).
13. To remove any impurities, the dialyzed sample was passed through a “Superdex 200 Increase” size-exclusion column (GE HealthCare, PA, USA) equilibrated with the BurrH solution. The size-exclusion run was performed using the BurrH solution and the absorbance at 280 nm was used to track BurrH.

#### Construction of p9344-BurrH

p9344-BurrH (GenBank MT081208) was created in two steps. First, an insert that included the Ampicillin resistance gene and plasmid origin was amplified by polymerase chain reaction from the pUC18 vector using primers:

5’-AAAAAAGAATTCCTTCCGCTTCCTCGCTCACTGACTC-3’

5’-AAAAAAGTCGACCTGCTCCCGGCATCCGCTTACAGAC-3’

The PCR product was cleaved by EcoRI and SalI, and ligated into the M13mp18 RF plasmid that was also cleaved by EcoRI and SalI. Next, the BurrH binding site was inserted into this construct. The 54-bp insert that comprises the BurrH binding site was ordered from IDT and inserted into the modified M13mp18 RF plasmid via SalI and HindIII restriction sites (Figure S1).

#### p9344-BurrH ssDNA Preparation

XL1-Blue MRF’ Kan was chemically transformed with p9344-BurrH plasmid. Next, the transformed XL1-Blue MRF’ Kan p9344-BurrH cells were mixed with molten LB top agar and spread onto an LB Agarose plate that was supplemented with 30 µg/ml Kanamycin. The plate was stored at 37°C overnight. The next morning, a plaque was picked and inoculated into 10 ml 2XYT media supplemented with 30 µg/ml Kanamycin, 100 µg/ml Ampicillin and 10 mM MgCl_2_. The inoculated culture was incubated overnight at 37°C with moderate shaking. On the next day, uninfected XL1-Blue MRF’ Kan culture in 100 ml 2xYT was supplemented with 30 µg/ml Kanamycin and 10 mM MgCl2 was grown at 37°C until the OD_600_ reached 0.4–0.6, and then 100 ml of that culture was inoculated with the overnight-grown XL1-Blue MRF’ culture producing p9344-BurrH phage. The 100-ml inoculated culture was incubated at 37°C with moderate shaking for 5 hours. p9344-BurrH phage was recovered and p9344-BurrH ssDNA was purified as described previously ^23^.

#### DNA Origami Designs and Preparation

p9344-BurrH scaffold routings and initial Goniometer designs were made using Cadnano2 ^24^. Optimization of staples (i.e. staple auto-breaking) and generation of pipetting instructions for the staple stocks were performed using a custom software toolkit (manuscript in preparation) developed in the Douglas Lab. Staple stocks for the goniometers were prepared using Labcyte Echo 550 (Labcyte, CA, USA). Folding reactions were performed using Bio-Rad MJ Research PTC-240 Tetrad thermal cycler. The temperature annealing ramp for the DNA Origami Goniometers was:

1. Incubate at 65°C for 00:10:00
2. Incubate at 60°C for 01:00:00 Decrease by 1.0°C every cycle
3. Cycle to step 2 for 20 more times

Folding was performed in 1XFOB20 (5 mM Tris-Base, 1 mM EDTA, 5 mM NaCl, 20 mM MgCl_2_ at pH 8.0). In the folding reaction, p9344-BurrH scaffold concentration was 20 nM and each staple concentration was 200 nM. The Goniometers were purified via PEG precipitation as described in ^25^. Briefly, 15% PEG-8000 solution (15% w/v PEG-8000, 5 mM Tris-Base, 1 mM EDTA, 500 mM NaCl, 20 mM MgCl_2_ at pH 8.0) were mixed with the folding reaction at a 1:1 ratio. The mix was centrifuged at 16,000× rcf for 25 minutes at room temperature. The supernatant was discarded and the pellet resuspended in 1XFOB20. PEG precipitation was repeated one more time and the final pellet was resuspended in 1XFOB20 and stored at 4°C.

#### Negative stain TEM experiments

Negative stain TEM experiments were performed on grids with different support surfaces for DNA-origami goniometer deposition. Prior to sample deposition and negative staining, ultrathin carbon coated quantifoil grids (Pacific grid tech, CA) were treated with a Pelco glow discharge unit (30 seconds hold/30 seconds glow discharge). For negative-stain grid preparation, 5 µl of 80 ng/ul DNA origami goniometer sample was deposited onto a glow-discharged ultra thin-carbon coated, graphene-oxide (made in-house) or amine-functionalized graphene-oxide grids (made in-house as described^15^). The sample was incubated on the grids for 1 minute. After incubation, the excess sample was wicked away using filter paper. Next, 10 µl of freshly prepared 2% Uranyl Formate (Electron Microscopy Sciences, PA) was applied to the grids and immediately wicked away using filter paper. A second round of 10 µl 2% Uranyl Formate was then applied onto the grids for 3 minutes before excess stain was again wicked away and the grid left to dry. Micrographs for the negatively stained grids were collected on Tecnai T12 (FEI, OR) at 30,000× magnification.

#### Cryo-EM Sample Preparation and Data Acquisition

For cryo-EM sample preparation, we used amine functionalized graphene-oxide grids prepared in-house as described previously^26^. Samples for cryo-EM were prepared by mixing 9 µl of 80 ng/µl DNA origami goniometer with 1µl of 2.5 µM BurrH in 1XFOB20 buffer supplemented with 1mM TCEP. Grids were prepared using Vitrobot (Thermo Fisher Scientific, MA) at 20°C. Prior to grid freezing in liquid ethane, the sample was incubated on the grid for 30 seconds and then blotted with a filter paper for 4 to 7 seconds. Micrographs were collected on a Talos Arctica microscope (Thermo Fisher Scientific, MA) operating at 200 kV with a K3 camera (Gatan, CA),at a nominal magnification of 22,000× corresponding to a physical pixel size of 1.82 Å. Total dose rate for imaging each hole is kept at 56 e/Å^2^.

#### Cryo-EM Image Processing and protein 3D reconstruction

Drift correction of the movie stacks was performed using MotionCor2 ^27^. The CTF parameter estimation of the drift corrected micrographs was performed using GCTF ^28^. Micrographs with ice contamination and with estimated resolution lower than 8Å were discarded from further analysis. Our data analysis pipeline for DNA origami goniometer classification included the following steps:

1. Goniometer particles were picked using Relion’s Laplacian of Gaussian Picking function ^16^.
2. Particles were classified into 2D classes for particle clean-up using Cryosparc ^17^. Bad classes were removed from further analysis (Fig. S6B).
3. Good classes were recentered and aligned to a single reference (Fig. S6C–D).
4. All regions of the goniometer except the bottom (tilt) barcode were subtracted and filled with gaussian noise centered at zero with a standard deviation of one (Fig. S6E).
5. Classifications of the subtracted goniometers that focused on the left and right portions of the bottom barcode region were performed using Relion’s 2D classification function ^16^. For the focused masked classifications, masks were created using EMAN2’s 2D mask drawing tool ^29^. Left and right bottom barcode classifications were performed successively until the classification results converged. Finally, classes that could not be recognized based on the barcode information were discarded (Fig. S6F–H).
6. Images with particles that were oriented with a flipped orientation relative to a chosen reference are flipped so that side (rotation) barcodes could be classified in a single classification run. The particles were aligned to a common reference and all regions of the goniometer except the side (rotation) barcodes were subtracted and filled with gaussian noise centered at zero with a standard deviation of one. Left and right side barcode classifications were performed successively until the classification results converge. Finally, classes that could not be recognized based on the barcode information were discarded. Side barcode classification was performed separately for each goniometer class (Fig. S6I–L).

After goniometers were separated based on the barcode bits, we moved onto data analysis steps that only involve the target protein:

1. Goniometers were aligned to a common reference so that target protein was at the center (Fig. S8A).
2. Protein particles were picked with box size as large as possible avoiding any signal from the goniometer (Fig. S8B).
3. Protein particles were classified using Relion’s 2D classification tool with the *psi* angle restrained to maintain protein orientations fixed with respect to Goniometers ^16^. Bad classes were removed from further processing (Fig. S8C).
4. Good classes were recentered keeping protein orientations fixed (Fig. S8D).
5. The initial 3D model was built using Relion’s 3D Initial Model function with additional arguments “`--sigma_tilt 10 --sigma_psi 10 --sigma_rot 15`” to keep the euler angles restrained around the *a priori* values ^16^, Fig. S8E).
6. All particles were aligned to the initial 3D model using Relion’s 3D Classification utility for single class classification with regularization parameter (T) set to 6 for stronger particle alignment and with additional arguments “`--sigma_tilt 10 --sigma_psi 10 --sigma_rot 15`” to keep the euler angles restrained around the *a priori* values ^16^, Fig. S8F).
7. Particles were classified into five 3D classes with regularization parameter set to 2 with additional arguments “`--sigma_tilt 10 --sigma_psi 10` “to keep the particle orientation restrained while allowing rotation angles to float (Fig. S8G).
8. 3D class with the best resolution, orientational, and positional accuracy, were picked and refined using Relion’s 3D refinement utility with additional arguments “`--sigma_tilt 10 --sigma_psi 10` “to keep the particle orientation restrained allowing rotation angles float ^16^, Fig. S8H).

In our data analysis pipeline, conversions between Cryosparc ^17^, Relion ^16^ and Cistem ^18^ data formats, data editing, particle subtractions and alignments were performed using custom Python scripts (https://github.com/douglaslab/cryoorigami). Cryo-EM data analysis was performed on AWS GPU instances p2.16xlarge and g3.16xlarge (Amazon, WA).

#### Model fitting into cryo-EM maps and model FSC calculations

The crystal structure of BurrH bound to DNA (PDB id: *4cja*) was fitted into 3D maps using UCSF Chimera’s rigid body fitting utility ^30^. FSC curves between the fitted *4cja* maps at 3.64Å resolution and the 3D refinement maps were calculated using EMAN2’s *e2proc3d.py* ^*29*^. Renders of the maps were generated using UCSF Chimera ^30^.

**Figure S1.**
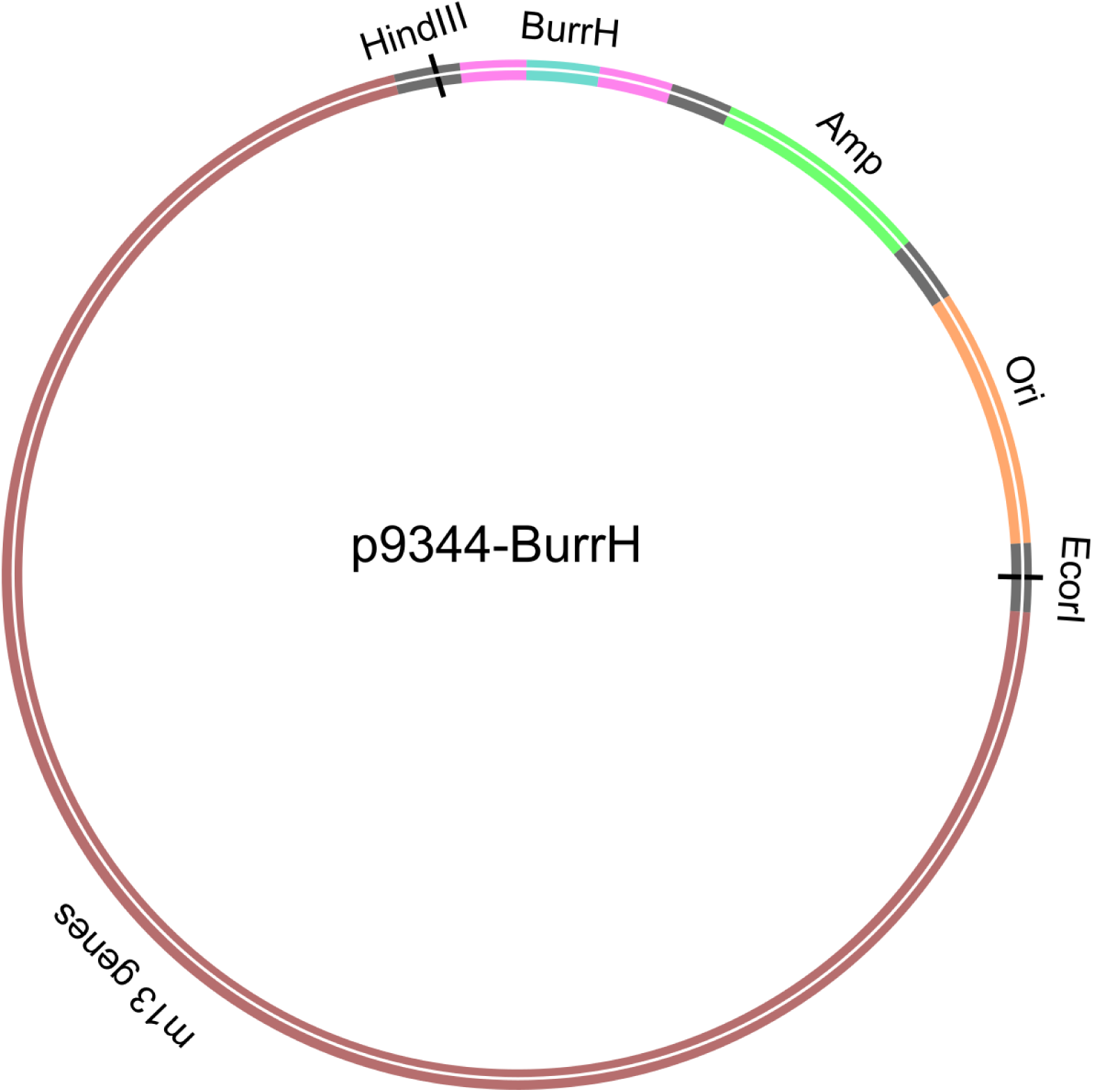
p9344-BurrH plasmid map. p9344-BurrH is created by inserting Ampicillin resistance gene (green), a plasmid origin of replication (orange) and BurrH binding site (green, magenta).

**Figure S2.**
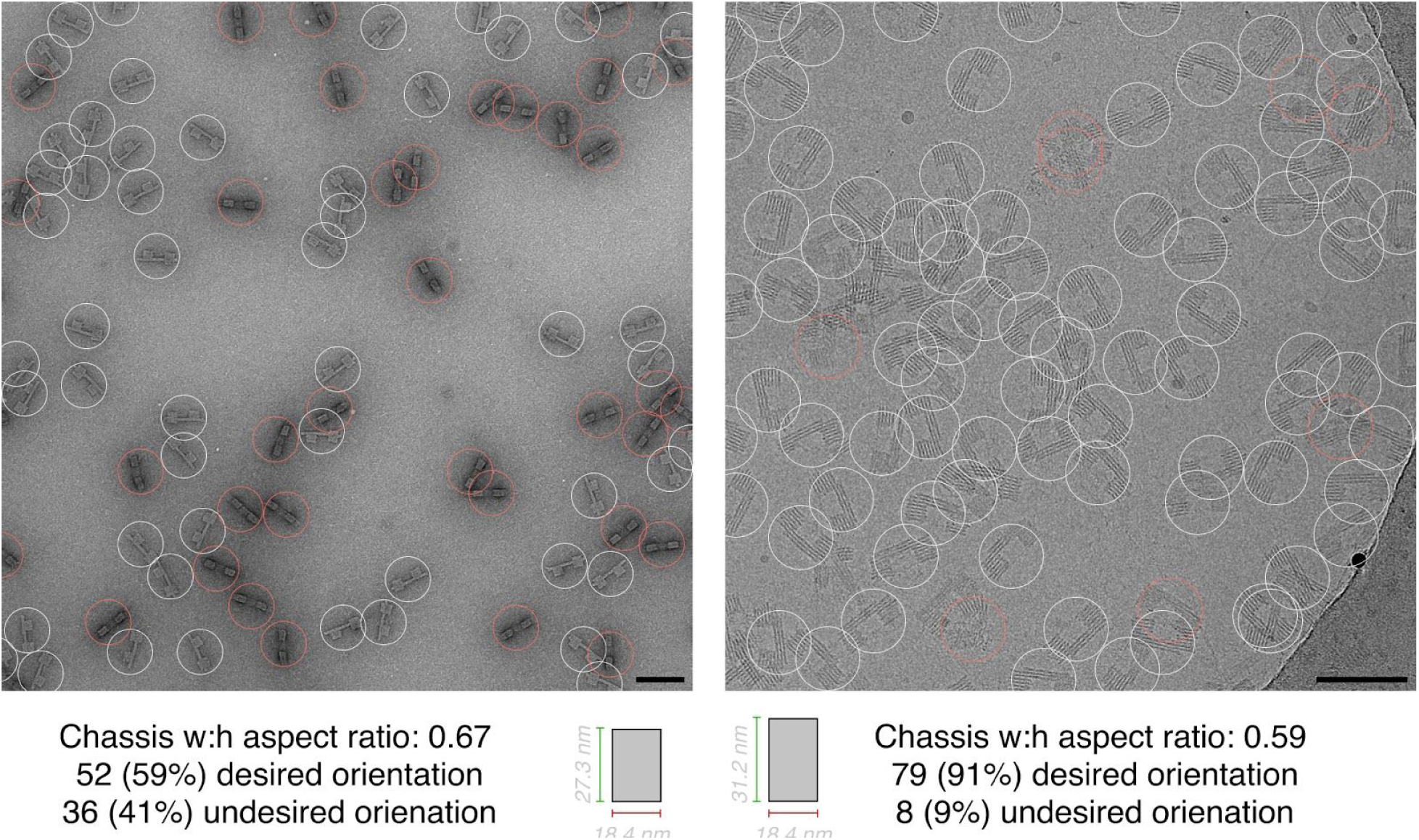
Comparison of grid adsorption orientation for two chassis aspect ratios. Representative micrographs of chasses with aspect ratios of 0.67 (left, negative stain) and 0.59 (right, cryo) were imaged and assessed by manual counting in. Scale bars: 100 nm.

**Figure S3.**
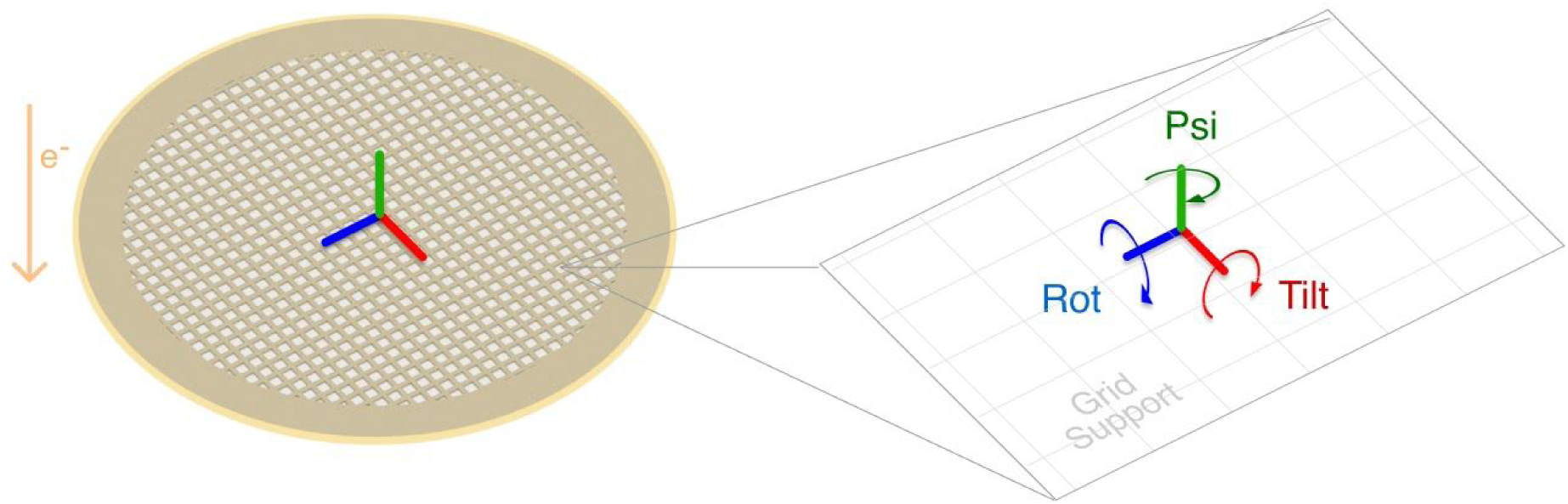
How we defined “tilt” and “rotation” angles for the molecular goniometers. Independent 2D views of a 3D object can be derived using only two orthogonal rotational transformations. In cryo-EM, the two orthogonal rotations can be referred to as tilt and rotation angles, respectively (Scheres 2012; Heymann et al. 2005). The reference coordinate system for the rotation operations can be chosen arbitrarily, and here we define the goniometer tilt angle as the angle between the stage DNA and the normal vector perpendicular to the goniometer face (the normal vector is parallel to the electron beam (Fig. 1A, orange line labeled e^−^) when the goniometer adsorbs in the desired face-up orientation. We define the goniometer rotation angle as the rotation angle with respect to the axis parallel to the DNA stage helical axis.

**Figure S4.**
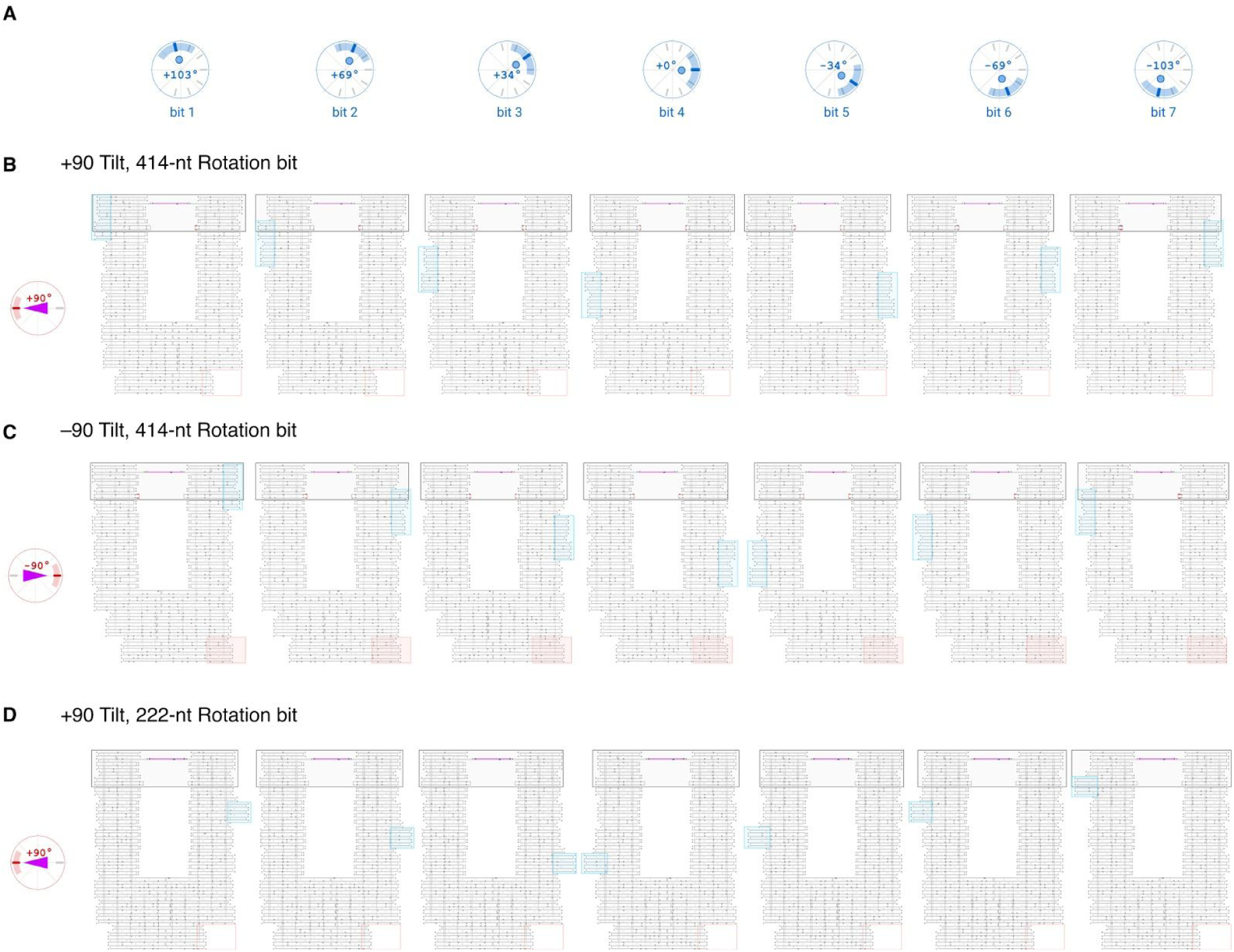
Cadnano design strand diagram schematics. (**A**) Bit and rotation angles for each column. (**B**) +90 tilt, 414-nt rotation bit designs. TILT bit is inactive (red outline in bottom right corner of each schematic). (**C**) –90 tilt, 414-nt rotation bit designs. TILT bit is active. (**D**) +90 tilt, 222-nt rotation bit designs. TILT bit is inactive.

**Figure S5.**
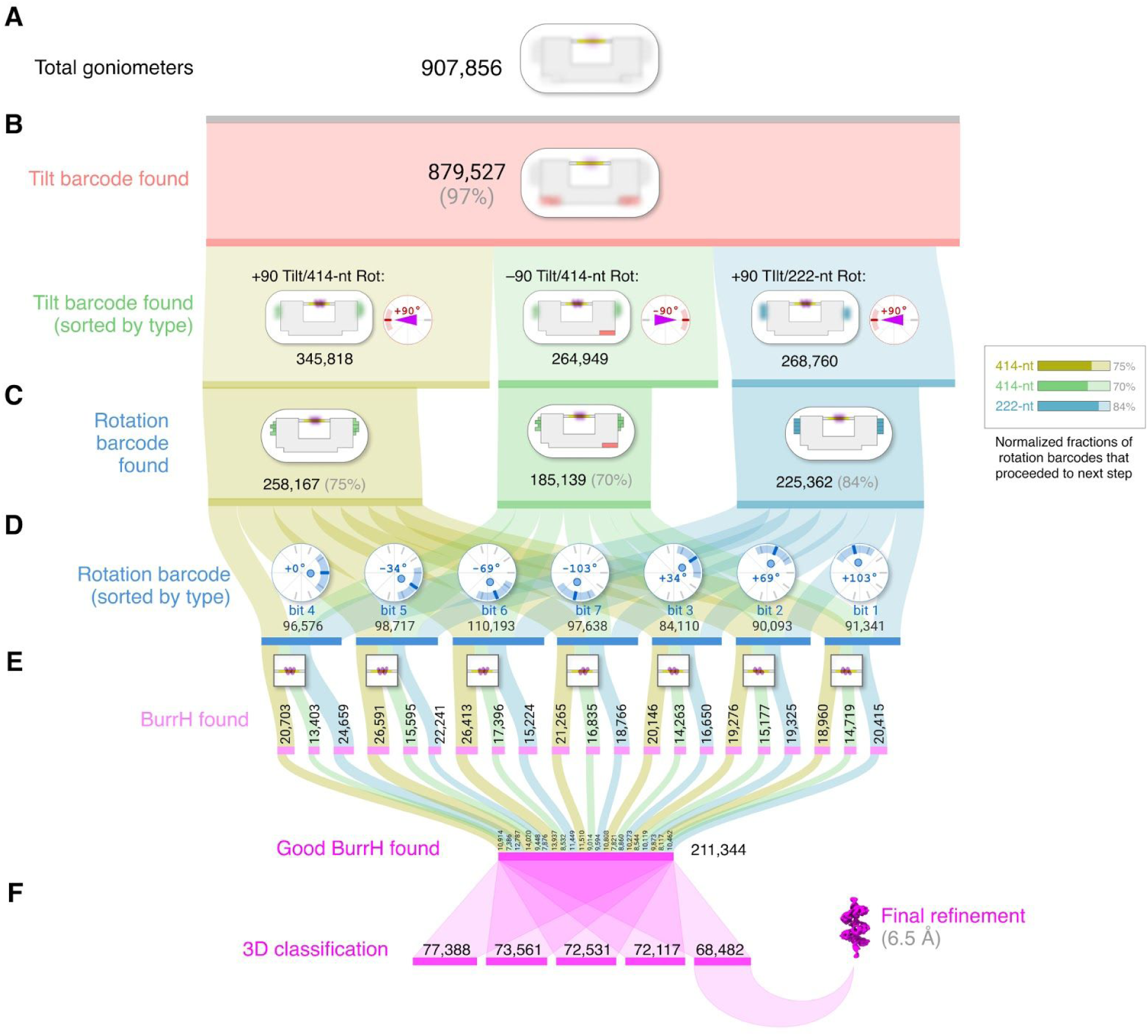
Sankey diagram for particle selection, classification, prior assignment, and 3D reconstruction. (**A**) “Total goniometers” is the count of particles successfully identified as a DNA origami goniometer. (**B**) “Tilt barcode found” means tilt barcode (Fig. 2B) was successfully classified. (**C**) “Rotation barcode found” means both tilt barcode and rotation barcode (Figs. 2B–D) were successfully classified. (**D**) Subclassification by rotation barcode for angle prior assignment. (**E**) “BurrH found” describes goniometers from (C) that contained a BurrH particle, and “Good” means the 2D BurrH subimage had a low background signal. (**F**) Good BurrH particles were used as input for 3D classification, and the highest-resolution class was used for final refinement.

**Figure S6.**
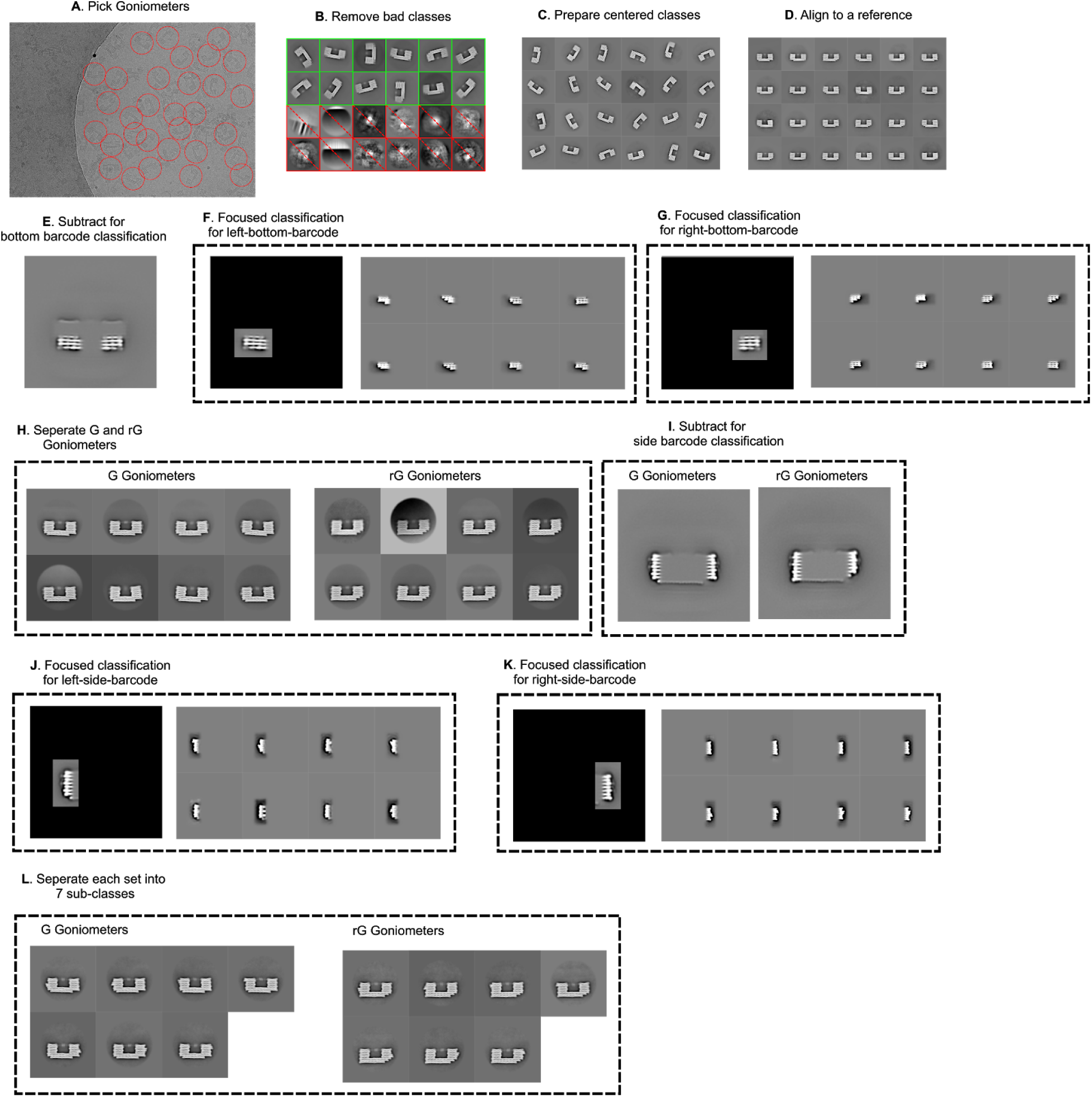
Cryo-EM data analysis workflow for DNA origami goniometer barcode classification. (**A**) Picking goniometers. (**B**) Removing bad goniometer classes. (**C**) Preparing well-centered classes. (**D**) Aligning goniometer classes to a reference. (**E**) Subtracting top portion of goniometers for bottom (tilt) barcode classification. (**F**) Focused classification on the left-bottom tilt barcode. (**G**) Focused classification on the right-bottom tilt barcode. **(H)** Separating +90° tilt and –90° tilt goniometers based on the barcode classifications. (**I**) Subtracting central portion of goniometers for rotation-barcode classification. (**J**) Focused classification on the left-side rotation barcode. (**K**) Focused classification on the right-side rotation barcode. (**L**) Separating goniometers into 7 sub-classes based on the rotation-barcode classifications (G = +90 tilt 414-nt rotation bit, rG = –90 tilt 414-nt rotation bit).

**Figure S7.**
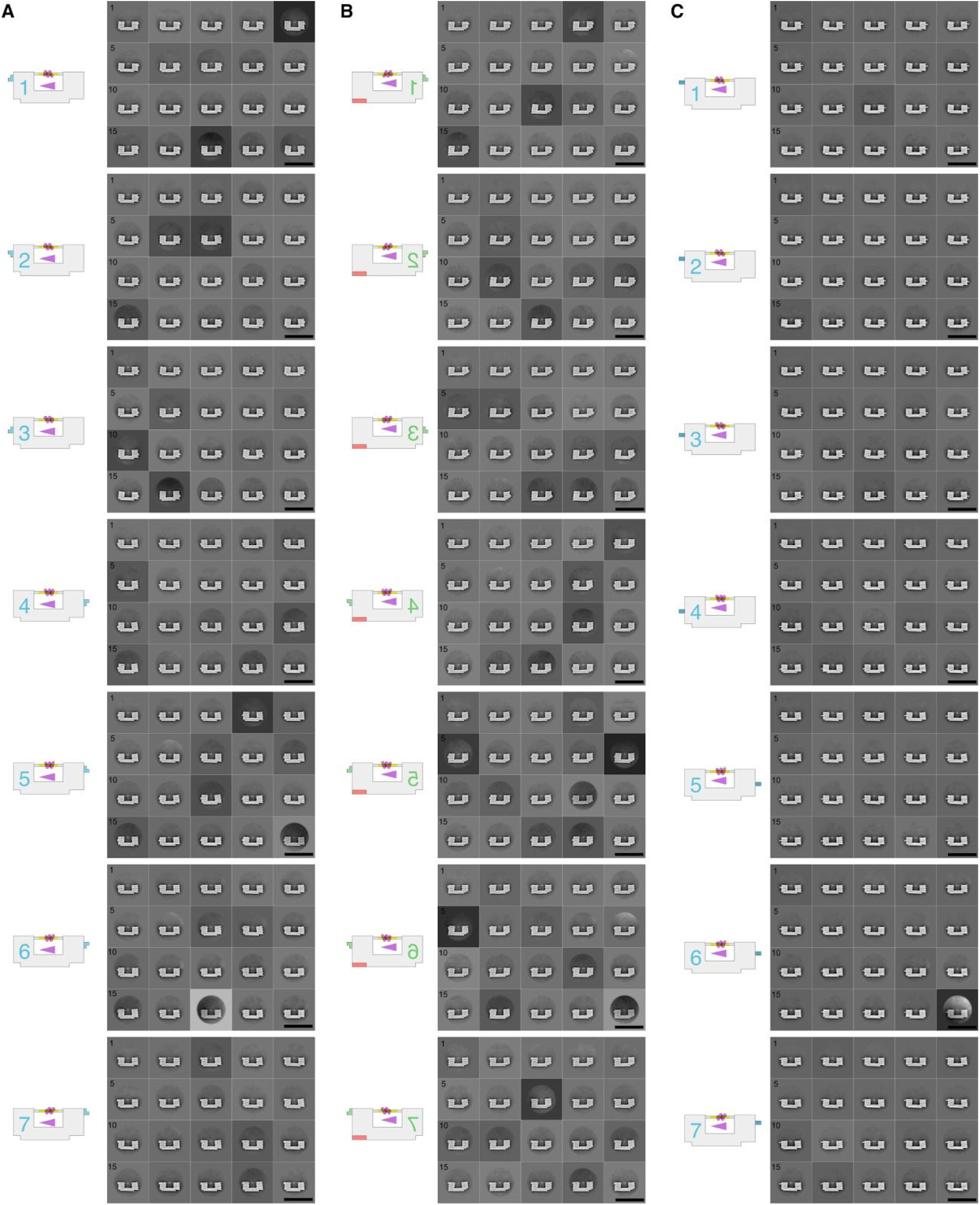
2D cryo-EM class averages goniometers with BurrH. (A) +90 tilt, 414-nt bit (B) –90 tilt, 414-nt bit (flipped), (**C**) +90 tilt, 222-nt bit. Scale bars: 100 nm.

**Figure S8.**
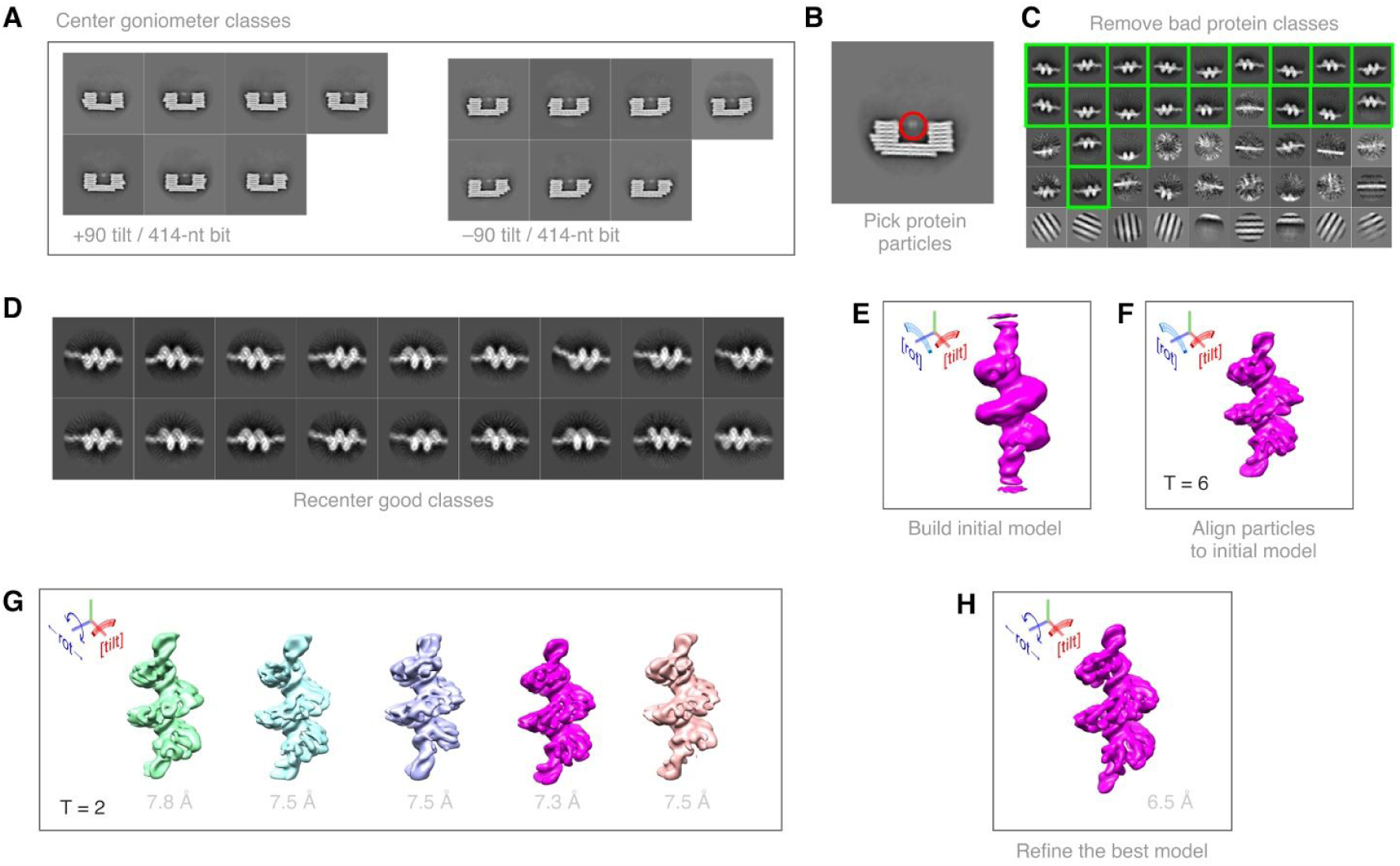
Cryo-EM data analysis workflow for BurrH 3D reconstruction and refinement. (**A**) Centering goniometer classes at tilt DNA. (**B**) Picking protein particles from the center of goniometers. (**C**) 2D classification of BurrH particles and removing “bad” classes. (**D**) Recentering “good” classes. (**E**) Building an initial model 3D model with tilt and rot angle constraints. (**F**) Aligning BurrH particles to initial model with tilt and rot angle constraints and regularization parameter, T, set to 6 for better alignment. (**G**) Classifying particles into five 3D classes with tilt angle constraint on and regularization parameter, T, set to 2 to minimize overfitting during classification. Best 3D class (magenta) is at 7.3 Å resolution. (**H**) Refining the best 3D class with only the tilt angle constraint. Estimated resolution of the refined map is 6.5 Å.

**Figure S9.**
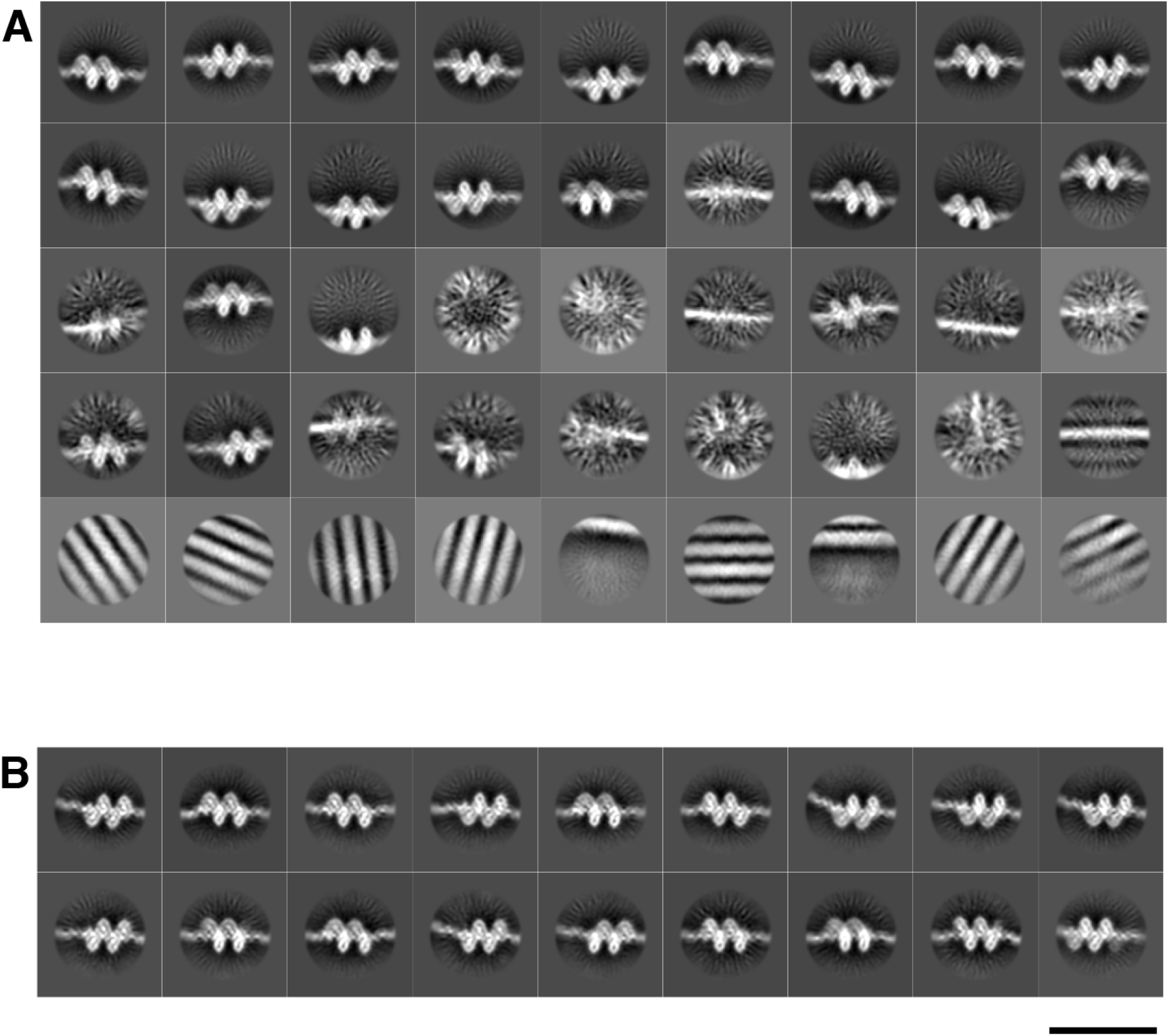
2D classification of goniometer tilt DNA with BurrH. **A.** Representative 2D classes of tilt DNA with BurrH after one round of 2D classification. Classes with high background signal, with empty tilt DNA and DNA origami signal are removed before further 2D classification of BurrH bound to tilt DNA. **B.** “Good” 2D classes retained for 3D reconstruction and refinement of BurrH. Scale bar is 20 nm.

**Figure S10.**
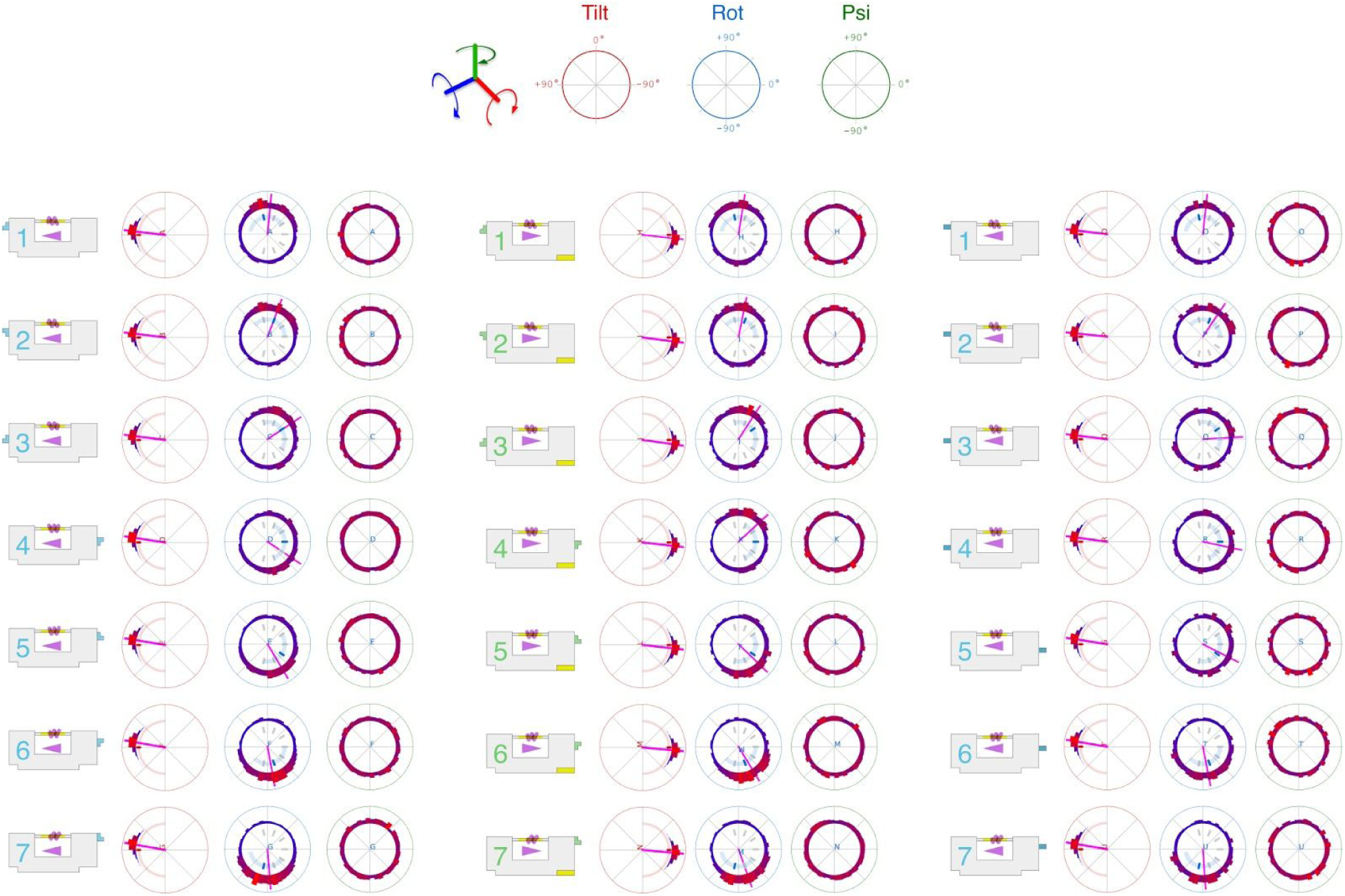
Measured tilt, rotation, and psi angle distributions for all goniometer designs. Polar plots and histograms follow the convention from Figure 3. The inner tick mark represents the goal angle, and the arc shows the bounds of the Gaussian angle prior. Average rotation angles are plotted as magenta lines for tilt and rotation plots.

